# Beyond Traditional Poincaré Analysis: Second-Order Plots Reveal Respiratory Effects in Heart Rate Variability

**DOI:** 10.1101/2025.08.13.670042

**Authors:** Mikhail A Lebedev, Alexandra S Medvedeva, Ksenia P Solovieva, Nadezhda M Starodubtseva, Anna V Makarova, Daria F Kleeva

**Affiliations:** Lomonosov Moscow State University, Moscow, Russia; Sechenov Institute of Evolutionary Physiology and Biochemistry of the Russian Academy of Sciences, St. Petersburg, Russia

**Keywords:** Heart rate variability, Poincare plot, cardiorespiratory, serial correlation, RR intervals

## Abstract

Heart rate variability (HRV) is a non-invasive biomarker of autonomic nervous system activity, commonly analyzed using a Poincaré plot. This plot visualizes correlations between successive heartbeats (RR_i_ vs. RR_i+1_) and quantifies autonomic regulation through SD1 and SD2 parameters. We introduce a second-order Poincaré plot, a natural extension that clarifies serial dependencies by plotting successive differences in RR intervals (ΔRR_i_ vs. ΔRR_i+1_). Applied to a PhysioNet dataset of 20 healthy individuals, this technique filtered out the slow HRV baseline of traditional elliptical plots to reveal distinct higher-order dynamics. These included ring-shaped structures indicating cardiorespiratory synchronization. A coupled-oscillator model, developed to simulate respiratory modulation, confirmed that these patterns are dictated by the respiratory frequency to heart rate ratio: slower breathing produces positive serial correlations in ΔRR, while faster breathing induces negative ones. By visualizing serial dependencies that conventional HRV metrics miss, the second-order Poincaré plot extends the classical analysis framework. This tool provides a refined method for uncovering subtle dynamical features in HRV across diverse physiological and clinical states.

**Highlights:**

- Second-order Poincaré plots, plotting successive differences of RR intervals (ΔRR_i_ vs. ΔRR_i+1_), extend traditional Poincaré analysis to reveal rapid HRV dynamics.
- In a dataset of 20 healthy individuals, second-order plots filtered out slow HRV components, highlighting respiratory modulation.
- Ring-shaped patterns in some participants indicated strong cardiorespiratory coupling, while others showed positive or negative serial correlations linked to breathing rate.
- A coupled-oscillator model confirmed that the ratio of respiratory to heart rate frequency determines serial correlation patterns.
- This method offers a novel tool for analyzing HRV dynamics, with potential applications in physiological and clinical research.

## 1. Introduction

Heart rate variability (HRV) captures the interplay between sympathetic and parasympathetic nervous system activity, providing insights into the heart’s adaptability to physiological and psychological demands (Shaffer & Ginsberg, 2017). As a non-invasive biomarker, HRV is widely used to evaluate cardiovascular health (Grässler et al., 2021), stress responses (Kim et al., 2018), and emotional regulation (Lane et al., 2009). Clinically, reduced HRV is consistently associated with adverse health outcomes (Lombardi, 2002), and its utility extends beyond cardiology to mental health research, where it serves as a valuable indicator (Kemp & Quintana, 2013). The rise of wearable technology has advanced HRV monitoring, enabling continuous, real-time data collection that enhances clinical diagnostics and personal wellness applications (Dalmeida & Masala, 2021; Li et al., 2023).

HRV analysis employs a range of methodologies, including time-domain and frequency-domain techniques, each offering distinct insights into autonomic nervous system regulation (Shaffer & Ginsberg, 2017). These methods assess autonomic function across various temporal scales, with the fastest fluctuations captured by analyzing consecutive heartbeat intervals, or RR intervals (Calabrese & Lambert-Lacroix, 2025; Tateno & Glass, 2001; Tsipouras et al., 2005). Key parameters for quantifying variability of consecutive RR intervals include the root mean square of successive differences (RMSSD) (Ciccone et al., 2017; Munoz et al., 2015) and the count of interval pairs differing by more than a specified threshold, such as 50 ms (NN50) (Mietus et al., 2002).

A Poincaré plot visualizes heart rate variability (HRV) by plotting each RR interval against the subsequent one (x = RR_i_, y = RR_i+1_). Initially applied by Raetz et al. (1991) to study RR interval correlations across sleep-wake states in cats, this method revealed distinct heartbeat patterns linked to physiological conditions. Since then, Poincaré plot analysis has become a standard tool for HRV assessment (Khandoker et al., 2013). Its characteristic elliptical point distribution yields two key parameters: SD1, which measures perpendicular dispersion and reflects parasympathetic-mediated short-term beat-to-beat variability, and SD2, which captures longitudinal dispersion and indicates combined sympathetic and parasympathetic influences on longer-term heart rate dynamics (Brennan et al., 2002; Tulppo et al., 1996; Voss et al., 2009). These metrics together offer a robust assessment of autonomic nervous system control over cardiac function.

This study advances Poincaré analysis by introducing second-order Poincaré plots, providing deeper insights into HRV dynamics. Unlike traditional first-order plots, which map consecutive RR intervals (RR_i_ vs. RR_i+1_), this method examines successive interval differences (ΔRR_i_ vs. ΔRR_i+1_). Applied to simultaneous electrocardiographic (ECG) and respiratory data from 20 healthy individuals, this approach revealed distinct, respiration-modulated HRV patterns unique to each subject. To elucidate these findings, we used a simple coupled oscillator model, where a rapid cardiac rhythm is dynamically modulated by a slower respiratory rhythm.

## 2. Methods

### 2.1. Data source and ethical approval

This study utilized data from the PhysioNet (Goldberger et al., 2000) dataset “Combined Measurement of ECG, Breathing, and Seismocardiograms Database”, contributed by Miguel Angel Garcia Gonzalez and Ariadna Argelagos Palou at Universitat Politècnica de Catalunya (García-González et al., 2013). The dataset includes recordings from 20 presumed healthy individuals, collected to address two main objectives: first, to examine whether subtle errors in RR interval detection across two ECG leads are influenced by respiratory activity; and second, to compare RR intervals from standard ECG measurements with surrogate estimates derived from seismocardiograms (SCG) to optimize SCG beat-detection algorithms. Data were acquired using a Biopac MP36 system (Santa Barbara, CA, USA), with two channels recording ECG (0.05–150 Hz bandwidth), a third channel capturing respiratory signals via a thoracic piezoresistive band (Biopac SS5LB sensor, 0.05–10 Hz bandwidth), and a fourth channel recording SCG signals using a triaxial accelerometer (LIS344ALH, ST Microelectronics, 0.5–100 Hz bandwidth). ECG electrodes (3M Red Dot 2560) with foam tape and conductive gel were used, and all signals were sampled at 5 kHz. Recordings were conducted with participants awake and motionless in a supine position on a standard bed. After sensor placement, a 5-min baseline recording was obtained, followed by approximately 50 min of classical music listening and a final 5-min rest period.

As the data were publicly available and de-identified, no additional ethical approval was required.

### 2.2. Signal processing

In this study, we analyzed the longest 50-minute music-listening epochs from the dataset, ensuring sufficient data for robust analysis. Signal processing and computations were performed using custom scripts in GNU Octave. The analysis commenced with the selection of the best-quality ECG channel for each participant (Fig. 1A). Following R-peak detection and timestamping, we processed sequences of four consecutive peaks (t_i_, t_i+1_, t_i+2_, t_i+3_) to compute successive RR intervals (RR_i_, RR_i+1_) and their differences (ΔRR_i_, ΔRR_i+1_). These metrics were then used to generate traditional Poincaré plots (RR_i_ vs. RR_i+1_; Fig. 1B) and novel second-order Poincaré plots (ΔRR_i_ vs. ΔRR_i+1_; Fig. 1C), enabling detailed exploration of heart rate variability dynamics. For the second-order Poincaré plots, we computed the serial correlation coefficient, defined as Pearson’s correlation between ΔRR_i_ and ΔRR_i+1_. RR intervals (ΔRR), computed as the difference between successive R-peak timestamps (t_i_, t_i+1_), were assigned a timestamp at the midpoint between them: (t_i_ + t_i+1_)/2. Similarly, the difference of successive RR intervals (ΔRR) was assigned a timestamp at the center weight of the three relevant R-peaks: (t_i_ + t_i+1_ + t_i+2_)/3. These discrete RR and ΔRR time series were then converted into continuous functions of time, RR(t) and ΔRR(t) using cubic spline interpolation (function interp1) and resampled at 5 kHz to match the original ECG and respiratory signals. This enabled direct comparison and the calculation of cross-spectral densities. For cross-spectral analysis (function cpsd), the continuous signals were downsampled to 5 Hz to focus on the respiratory frequency band (~0.3 Hz). Separately, to analyze the heart rate near ~1 Hz, ECG was downsampled to 50 Hz before computing its power spectral density.

**Figure 1.**
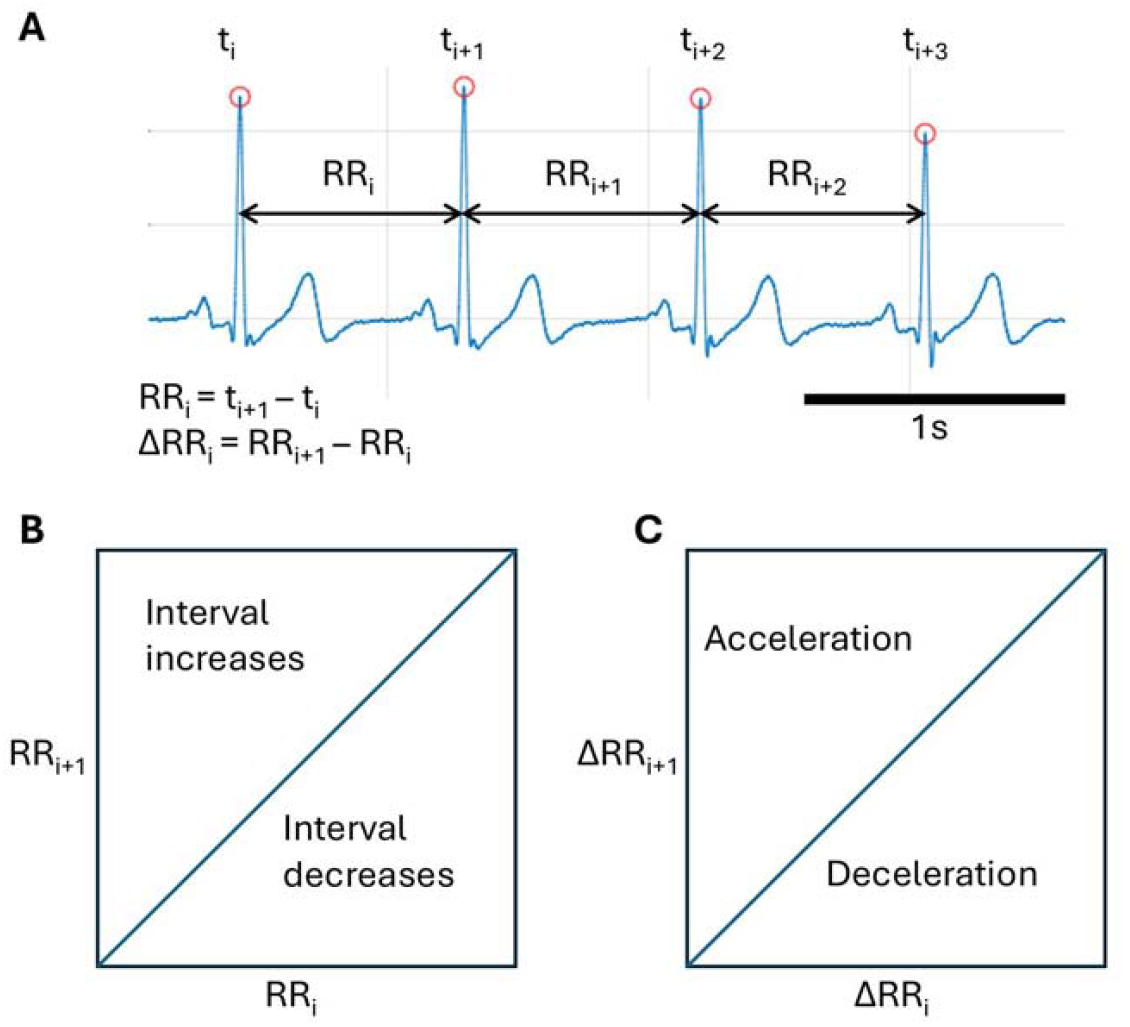
Construction of the first-order and second-order Poincaré plots. A: ECG trace with detected R peaks marked by red circles. Four consecutive peaks are timestamped, and their occurrence times are used to compute RR intervals and successive differences (ΔRR). B: First-order Poincaré plot with RR_i_ on the x-axis and RR_i+1_ on the y-axis. Points above the line y = x indicate RR interval lengthening, while points below the line indicate shortening. C: Second-order Poincaré plot with ΔRR_i_ on the x-axis and ΔRR_i+1_ on the y-axis. Points above the line y = x correspond to RR interval acceleration (ΔR increase), while points below the line indicate deceleration (ΔR decrease).

### 2.3. Coupled-oscillator model

We developed a simple coupled-oscillators model to simulate respiratory modulation of heart rate. The cardiac oscillator was implemented as an integrate-and-fire process. Its voltage rose at a rate equal to the inverse of the desired heart rate, and a spike was generated once it reached a fixed threshold. This baseline threshold was calibrated to produce the target heart rate in the absence of other inputs. To simulate physiological variability, we added noise to the threshold and a sinusoidal modulation at the respiratory frequency. As the model was designed for illustrative purposes, its parameter space was not exhaustively explored.

### 2.4. Statistical analysis

Pearson’s correlation coefficient, r, was calculated to evaluate serial correlations in second-order Poincaré plots, specifically between ΔRR_i_ and ΔRR_i+1_. The significance of r was tested using a t-test. Linear regression was used to analyze the relationship between the modulation frequency-to-heart rate ratio and the ΔRR serial correlation coefficient, r. The model, y = ax +b, where a is the slope and b is the intercept, was fitted using the least-squares method. A t-test assessed the significance for this regression. The analyses were performed in GNU Octave.

## 3. Results

Figure 2 presents the analysis for three participants from the dataset. A representative 5-minute interval is shown, taken from the 50-minute recording session. For each participant, recordings of respiration are shown and the corresponding values of RR and ΔRR as a function of time (Fig. 2A, C, E). Notably, participant 9 had an unusually low respiratory frequency (approximately 4 breaths per minute), visibly correlated with both RR and ΔRR (Fig. 2C). By contrast, in participants 3 (Fig. 2A) and 19 (Fig. 2E), breath frequencies were approximately 17 and 20 breaths per minute. In all participants, the plots for RR contain very slow heart rate fluctuations (0.01 – 0.1 Hz), which are filtered out when transitioning to ΔRR. In the classical Poincaré plots (Fig. 2B, D, F, top), a characteristic ellipse is present which corresponds to these slow fluctuations, with its major axis aligned along the line y = x. The second-order Poincaré plots (Fig. 2B, D, F, bottom) show additional patterns not evident in the first-order plots. Notably, the second-order plot for participant 3 (Fig. 2B, bottom) very clearly exhibits a ring-like structure, and a ring pattern can be also noticed for participant 19 (Fig. 2F, bottom). This feature is absent in participant 9 (Fig. 2D, bottom). The coefficient of serial correlation, r, is indicated for each second-order Poincaré plot: −0.03, 0.48, and −0.34, for participants 3, 9 and 19, respectively. These values match the orientation of the scatter patterns: for participant 9 (Fig. 2D, bottom), the points cluster along the line y = x (i.e. positive serial correlation) whereas for participant 19 (Fig. 2F, bottom) the point distribution is perpendicular to the line y = x (i.e. negative serial correlation). Importantly, none of these distinct patterns could have been anticipated from the first-order Poincaré plots alone.

**Figure 2.**
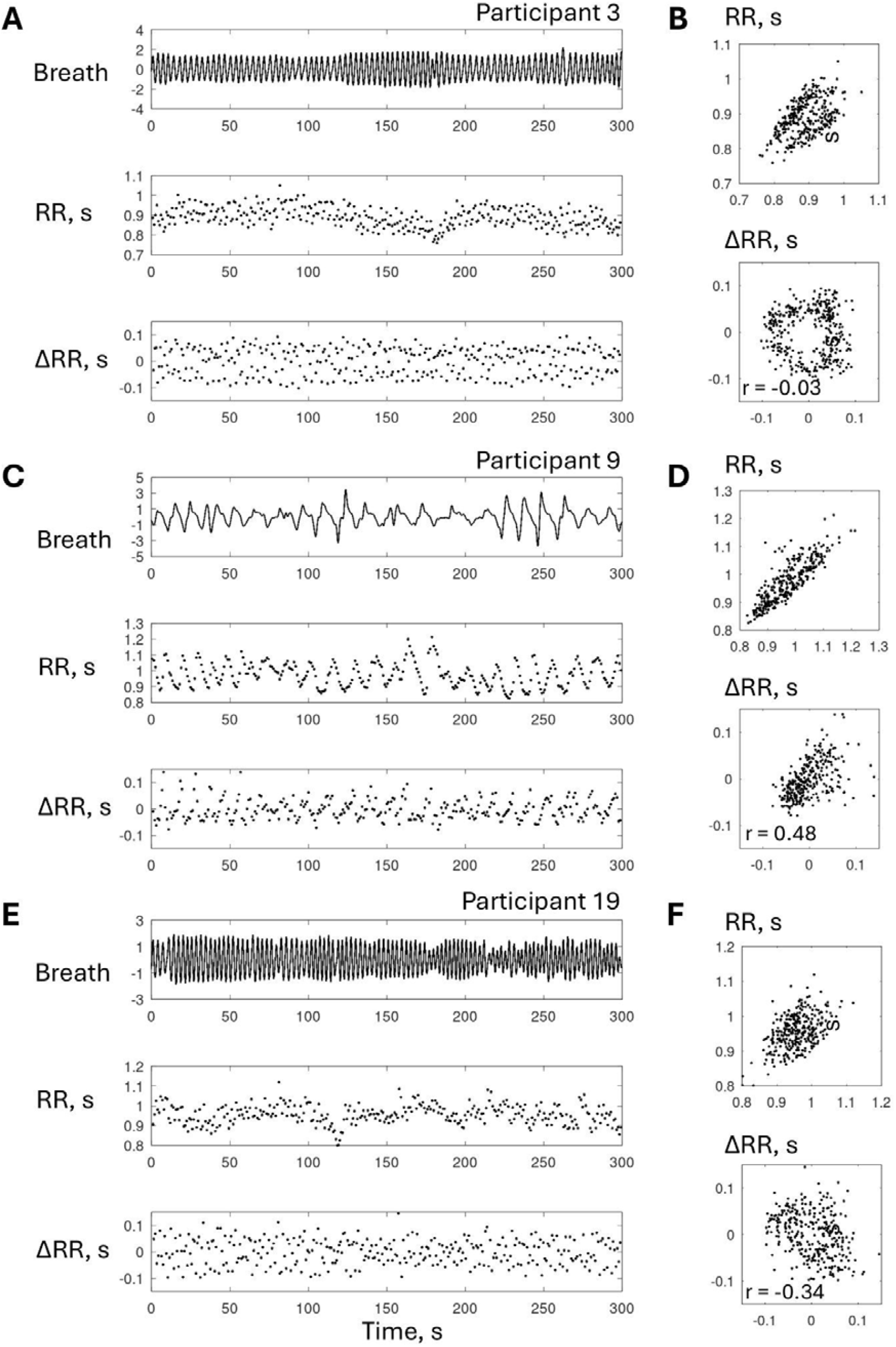
Time course of heartbeat intervals (RR) and their changes (ΔRR), along with first-order and second-order Poincaré plots for the same data. A, C, E: Respiration (top), RR (middle) and ΔRR (bottom) plotted over time for participants 3, 9, and 19, respectively. B, D, F: First-order (top) and second-order (bottom) Poincaré plots for the corresponding participants.

The ring-like pattern observed in the second-order Poincaré plot for participants 3 and 19, as well as in five other participants upon visual inspection, suggested that the respiratory cycle might be the primary driver of this effect. To investigate this possibility, we examined the modulations of RR intervals and their changes, ΔRR, in relation to recorded breath patterns. Figure 3 presents the analysis for the same participants as in Fig.2, with respiratory recordings aligned to peak inhalation at the top of panels A, B, and C. The corresponding traces for RR intervals and ΔRR are displayed in the middle and bottom sections of these panels, respectively. To facilitate comparison, the discrete data points derived from R peaks were converted into a continuous representation through interpolation. Breath cycles are clearly correlated with the modulations of RR and ΔRR in all three participants. Additionally, panels D-F in Fig. 3 illustrate the second-order Poincaré plots as three-dimensional visualizations, with time represented along the z-axis. The plots form spirals for participants 3 (Fig. 3D) and 19 (Fig. 3F), with each full rotation corresponding to a complete respiratory cycle. No such spirals are visible for participant 9 (Fig. 3E), which corresponds to the absence of the ring-like pattern for this participant (Fig. 2D, bottom). To confirm the correlation of ΔRR modulations with breathing, cross-spectra (black lines) were computed for respiration versus ΔRR, where the latter was treated as a continuous variable. The spectra for both respiration (green lines) and ΔRR (magenta lines) were also calculated. For participants 3, 9, and 19, this approach clearly identified spectral peaks corresponding to respiratory cycles (Fig. 3G-I). Spectral analyses were similarly performed on ECG records, revealing peaks associated with heart rate at around 1 Hz (Fig. 3J-L).

**Figure 3.**
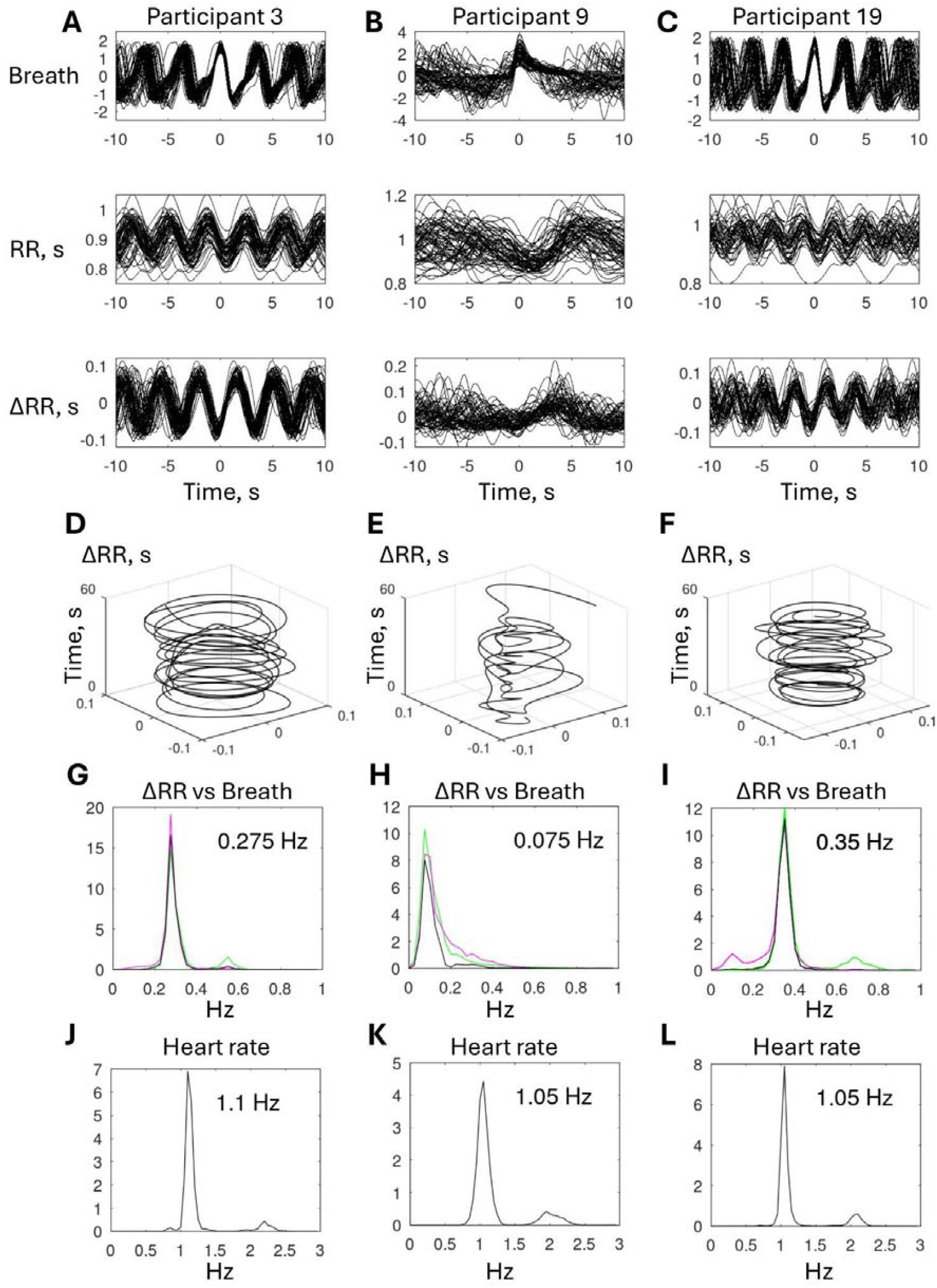
Comparison of respiratory cycles with modulations of RR intervals and ΔRR in participants 3, 9, and 19 (the same as in Fig. 2). RR and ΔRR values were interpolated to generate continuous traces. A-C: Respiratory traces (top) aligned to peak inhalation, with corresponding RR (middle) and ΔRR (bottom) modulations. D-F: Three-dimensional representations of the second-order Poincaré plots, with time as the third axis. Spiral patterns reflect respiratory cycles in D and F. G-I: Cross-spectral analysis of respiratory-related modulations. Black lines represent cross-spectra (ΔRR vs. respiration), while colored lines denote spectra for respiration (green) and ΔRR (magenta). Frequencies corresponding to the peaks of ΔRR–respiration cross-spectra are indicated. J-L: ECG spectra, with peaks corresponding to heart rate.

Additional examples of the second-order Poincaré plots and the corresponding spectra are depicted in Figs 4, 5, and 6, corresponding to ring patterns, positive serial correlation, and negative serial correlation, respectively. In these illustrated cases (and the other cases that are not shown), peaks are present in the cross-spectra for ΔRR versus respiration at 0.2-0.4 Hz, which reflect cardiorespiratory modulations, and the frequency of 0.075 Hz for subject 9 is unusually low (Fig. 3H). Peaks at the same frequencies are present in the spectra for ΔRR and respiration. In one illustrated case (Fig. 6B), low-frequency drifts are noticeable in the respiratory spectrum, reflecting imperfections of the recordings. Additionally, respiratory spectra are sometimes broad (Figs 5E and 6E) because of the variations in breath frequency. Additionally, ΔRR autospectra in some cases exhibit modulatory frequencies higher than the breath frequency (Figs 5E and 6E). While a detailed analysis of these cases is beyond the scope of this study, they may warrant further investigation, particularly in clinical applications.

**Figure 4.**
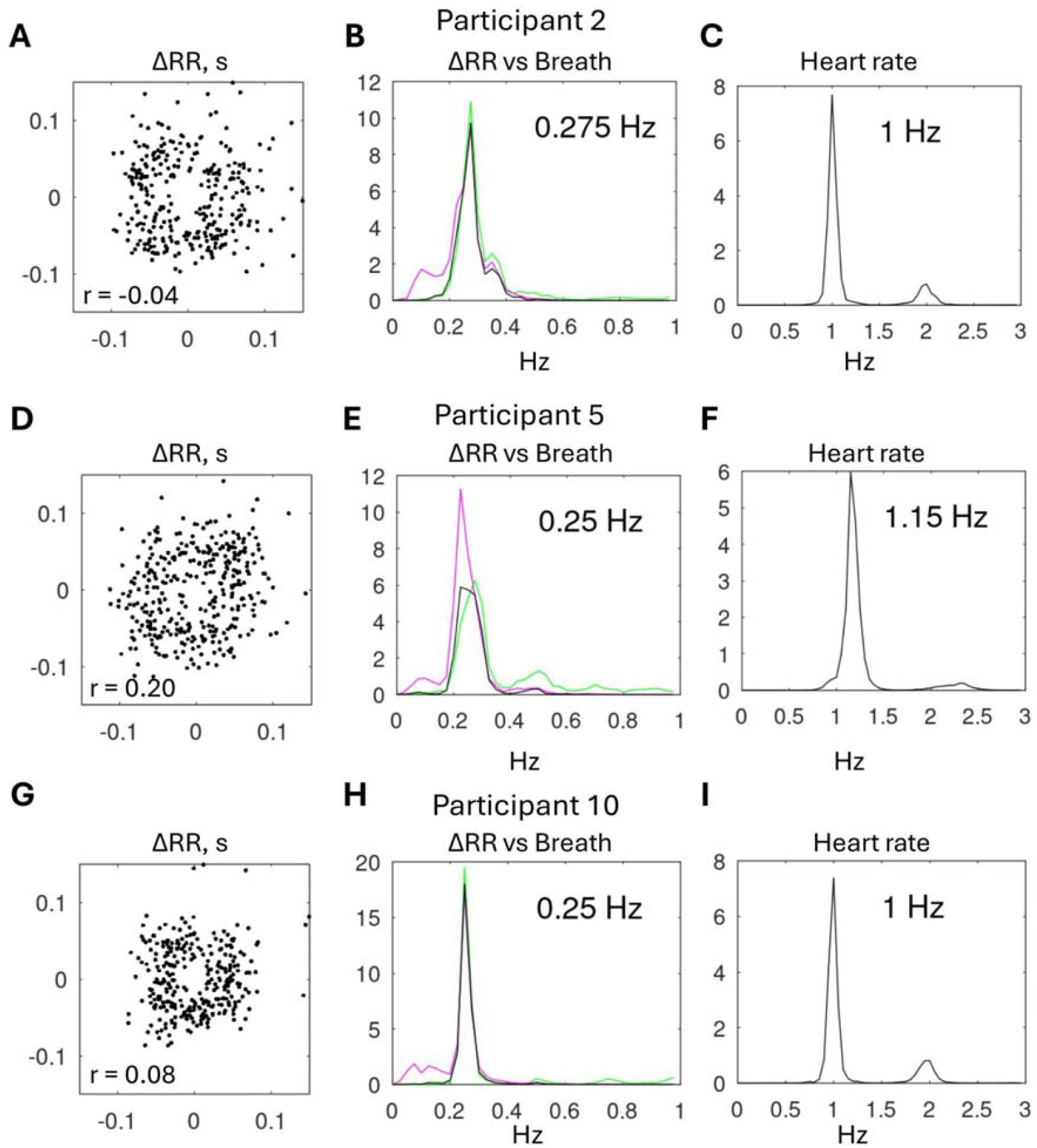
Examples of ring patterns in the second-order Poincaré plots and the corresponding spectra and cross-spectra. Results are shown for 5 minutes of recordings in participants 2, 5, and 10. A, D, G: Second-order Poincaré plots, with the coefficient of serial correlation, r, indicated. B, E, H: Cross-spectral analysis of respiratory-related modulations. Black lines represent cross-spectra (ΔRR vs. respiration), while colored lines denote spectra for respiration (green) and ΔRR (magenta). Frequencies corresponding to the peaks of ΔRR–respiration cross-spectra are indicated. C, F, I: ECG spectra, with peaks corresponding to heart rate.

**Figure 5.**
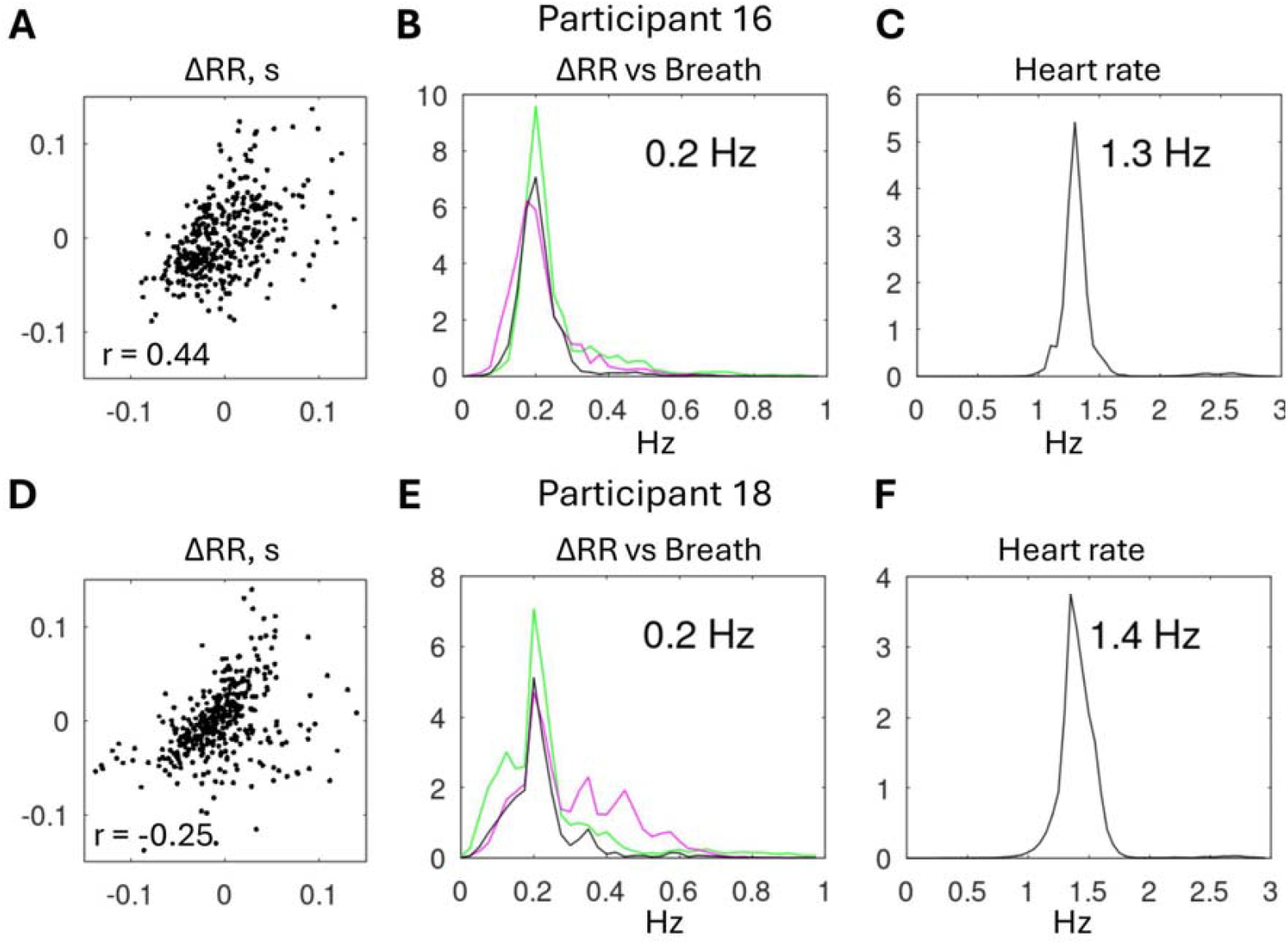
Second-order Poincaré plots and spectral analysis of heart rate versus respiration for two participants (16 and 18) with positive serial correlation for ΔRR. Conventions as in Fig. 4.

**Figure 6.**
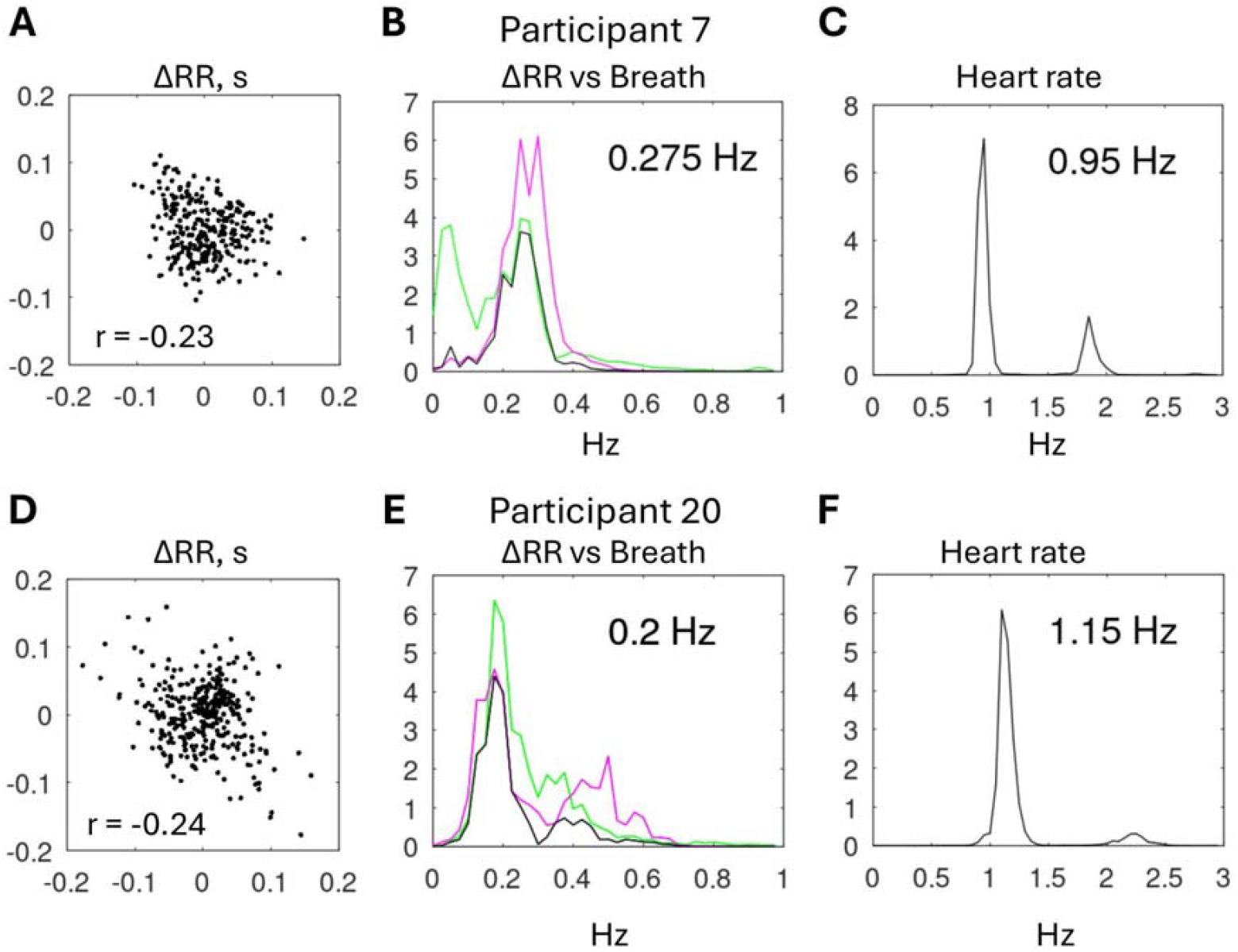
Second-order Poincaré plots and spectral analysis of heart rate versus respiration for two participants (7 and 20) with negative serial correlation for ΔRR. Conventions as in Fig. 4.

To summarize the results obtained thus far, we found that second-order Poincaré plots provided additional insights compared to first-order plots. These plots highlighted rapid modulations of RR intervals, contrasting with the slow modulations captured by the first-order plot ellipse. In some cases, they also revealed ring-like patterns associated with a strong correlation between ΔRR and breath cycles. (But a strong correlation did not necessarily translate to a ring pattern as shown in Figs 2C,D and 3B,E,H.) Additionally, second-order Poincaré plots exposed either positive or negative serial correlations which could not be predicted from the first-order plots.

We analyzed the individual variability in serial correlation in more detail. First, we noticed a trend for the lower breath rates to coincide with positive serial correlation and for the higher breath rates to align with negative serial correlation. Thus, participant 9 exhibited positive serial correlation (r = 0.48; Fig. 2D) alongside a very low breath rate (0.075 Hz, Fig. 3H), whereas participant 19 showed negative serial correlation (r = −0.34; Fig. 2F) with a higher breath rate (0.35 Hz; Fig. 3I). Second, we noticed that the heart rate also contributed to the serial correlation patterns. Thus, participants 16, and 18 displayed positive serial correlations with a low breath-ΔRR coupling frequency of 0.2 Hz and a somewhat elevated heart rate of 1.3-1.4 Hz (Fig.5).

Next, we considered the interaction between a high-frequency cardiac oscillator (generating pulses roughly every 1 s) and a low-frequency respiratory oscillator (whose cycle spans approximately 3 heartbeats), where altering the rate of either oscillator changes the number of RR intervals within each respiratory cycle. If the respiratory rate slows—and/or the heart rate increases—more RR intervals will fit into a single breath cycle. In such cases, ΔRR tends to follow a pattern where it increases (or decreases) progressively over the course of the respiratory cycle, leading to a positive serial correlation in ΔRR. Conversely, if the respiratory rate increases—and/or the heart rate decreases—fewer RR intervals occur within each breath cycle. As a result, the respiratory cycle becomes too short to sustain a consistent directional change in ΔRR, causing fluctuations that instead produce a negative serial correlation.

To verify whether this mechanism applied to our data, we examined the relationship between the breath-to-heart rate ratio (a measure proportional to the number of RR intervals per respiratory cycle) and the ΔRR serial correlation coefficient, as shown in Fig. 7. The figure presents two variants of this analysis. Figure 7A uses the respiratory frequency derived from cross-spectral analysis (representing the frequency at which respiration modulates ΔRR), while Figure 7B uses the dominant modulation frequency obtained from the ΔRR spectrum. As noted earlier, these frequencies typically coincided when respiratory modulation of heart rate was strong. However, discrepancies arose in certain cases due to additional non-respiratory heart rate modulations (e.g., Figs 5E and 6E) or questionable respiratory signal quality (e.g. Fig. 6B). This justified the inclusion of Fig. 7B’s analysis, which becomes particularly relevant when respiratory recordings are unavailable. Additionally, with long analysis epochs (such as 5 minutes in Figs 2-6), nonstationarities could occur in both breath and heart patterns, altering the results of these analyses. Therefore, for both analyses, the 50-min recordings were segmented into 100-second epochs (yielding 60 epochs per participant). Regression analysis confirmed our hypothesis in both cases, though the fit was stronger when using the modulation frequency derived solely from the ΔRR spectrum.

**Figure 7.**
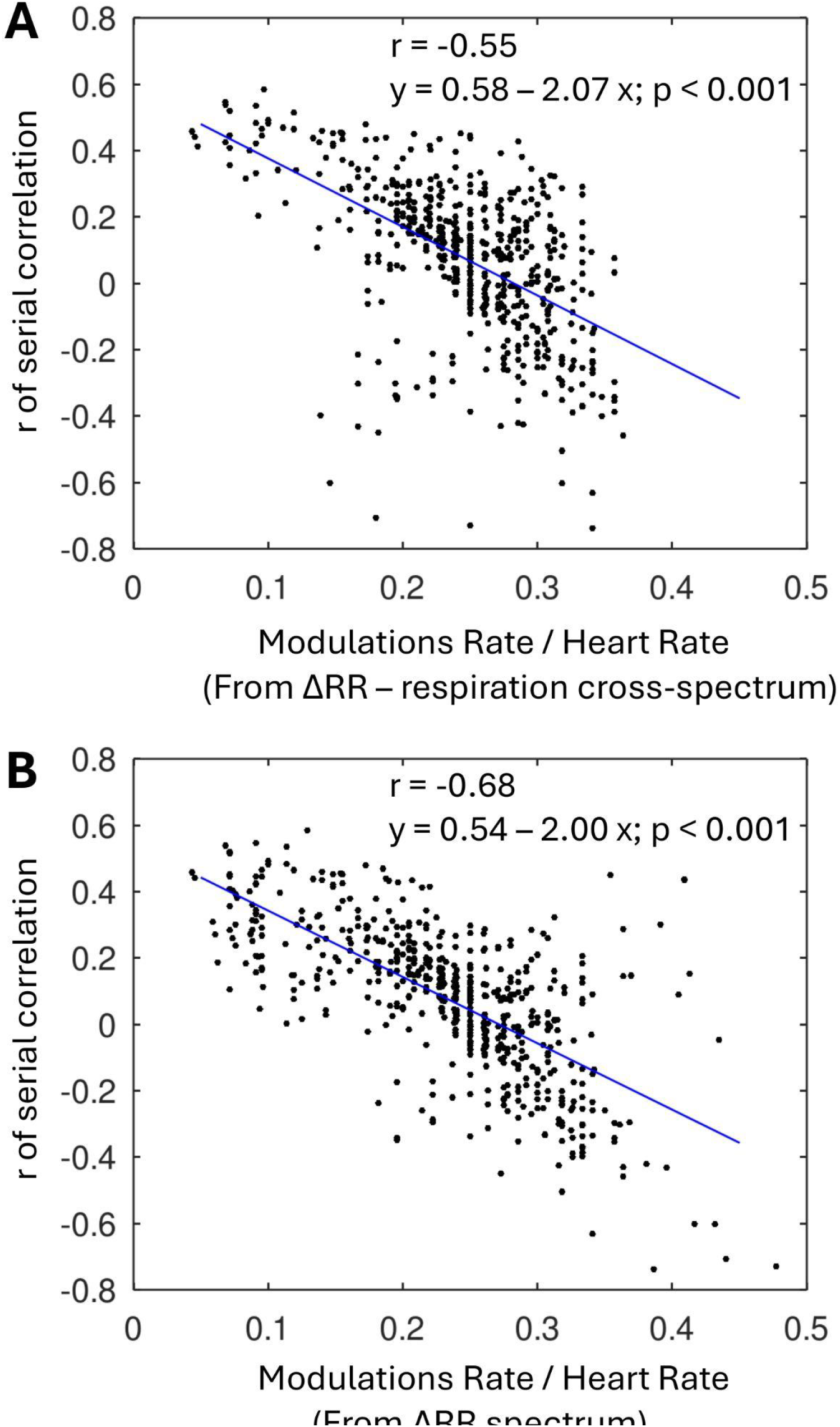
Relationship between modulation frequency-to-heart rate ratio (x-axis) and ΔRR serial correlation coefficient (y-axis). Each data point represents a 100-s epoch taken from a 50-min recording session. Solid lines show linear regression fits with corresponding equations (y = ax + b), correlation coefficients (r), and p-values indicated. A: Modulation frequency obtained from cross-spectral analysis of respiration and ΔRR. B: Modulation frequency obtained from spectral analysis of ΔRR alone.

To further verify the proposed mechanism of negative versus serial correlation in ΔRR for different breathing and heart rates, we developed a simple model of the coupling between an approximately 1 Hz cardiac oscillator with an approximately 0.3 Hz respiratory oscillator (Fig. 8). The cardiac oscillator was modeled using an integrate-and-fire mechanism with a constant rate of rise but a randomly fluctuating starting point (Fig. 8A). An impulse was generated when the integrated rate reached a threshold that was sinusoidally modulated according to the breathing rate. The model successfully reproduced the expected relationship between the ratio of respiration rate to heart rate, which was accurately described by a linear fit (Fig. 8B). Figure 8C displays second-order Poincaré plots for different model parameters, illustrating a transition from positive to negative serial correlation as breathing rate increased and/or heart rate decreased. Intermediate parameter settings also revealed distinct ring-like patterns, further supporting the model’s ability to capture the observed dynamics.

**Figure 8.**
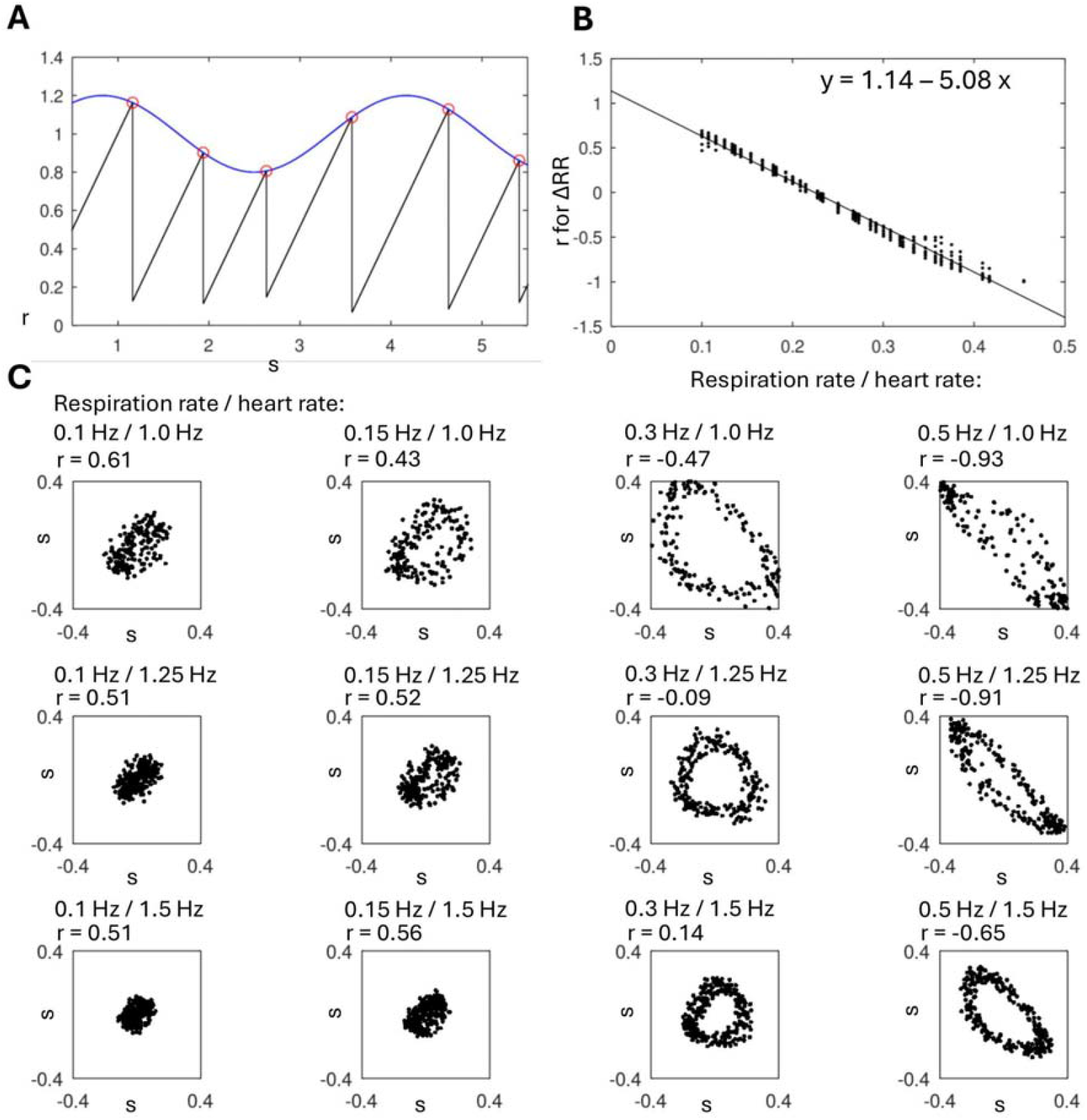
A simple model of coupling between a fast cardiac oscillator (~1 Hz) and a slow respiratory oscillator (~0.3 Hz). A: The cardiac oscillator was modeled as an integrate-and-fire pacemaker with a constant rate of rise but a randomly varying starting point. Impulses were generated when the integrated signal reached a threshold, which was sinusoidally modulated by respiratory activity. B: Linear regression of the respiration-to-heart-rate ratio against the serial correlation coefficient of ΔRR. C: Second-order Poincaré plots illustrating the shift from positive to negative serial correlation as breathing rate increased and/or heart rate decreased, with ring-like patterns emerging at intermediate parameter values.

## 4. Discussion

The present findings build upon the classical Poincaré analysis of HRV, as established in prior research (Brennan et al. 2002; Kamen et al. 1996; Khandoker et al. 2013; Raetz et al. 1991; Tulppo et al. 1996; Voss et al. 2009), by introducing a second-order Poincaré plot in which ΔRR_i_ is plotted against ΔRR_i+1_. This extension aligns naturally with the existing framework, given that interpretations of Poincaré plots are fundamentally rooted in the statistical properties of ΔRR. The visualization and quantification of serial dependencies in ΔRR thus provide a coherent and meaningful advancement of the methodology.

In the present study, the introduction of the second-order Poincaré plot improved the analysis of cardiorespiratory coupling—the physiological interplay between breathing rhythms and heart rate modulation. Respiration influences cardiac activity through well-established neural mechanisms, particularly respiratory sinus arrhythmia, characterized by heart rate acceleration during inspiration and deceleration during expiration (Berntson et al., 1993; Larsen et al., 2010; Yasuma & Hayano, 2004). Analysis of individual data revealed distinct coupling patterns, with some participants exhibiting pronounced ring-like structures in the second-order Poincaré plots, while others displayed varying degrees of serial correlation in ΔRR intervals. Cross-spectral analysis confirmed respiratory modulation in all cases. We proposed that breathing frequency relative to heart rate determines the observed individual differences: slower respiration produces sustained directional changes in successive RR intervals, yielding positive serial correlations, whereas faster breathing induces more frequent reversals in interval dynamics, leading to negative correlations. This hypothesis was tested using a simple integrate-and-fire oscillator model, where respiration modulated threshold crossings. This model supported the notion that respiratory frequency shaped the temporal structure of HRV, as captured by second-order Poincaré analysis.

Here we analyzed a single data set—the recordings from healthy individuals lying quietly in bed and listening to music. Under these conditions, respiratory modulation of heart rate is likely to be especially prominent. It would be informative to extend our analysis to other conditions in which first-order Poincaré analysis has been applied previously, for example heart-rate recordings during different breathing regimes (Guzik et al. 2005; 2007), during exercise (Mourot et al. 2004), and in studies of autonomic influences such as head tilts and pharmacological agents (Kamen et al. 1996; Karmakar et al. 2011). It would also be of interest to determine whether second-order Poincaré plots offer additional utility for clinical cases like arrhythmias, where first-order analysis has already been used (Zhang et al. 2015).

In conclusion, our results underscore the value of second-order Poincaré analysis for uncovering dynamical features that conventional HRV metrics may miss. Future research should examine whether similar patterns occur in other physiological oscillator systems and how these dynamics change across different physiological and cognitive conditions, disease states, or pharmacological interventions. The potential clinical applications warrant further exploration, particularly for conditions involving autonomic dysfunction or impaired cardiorespiratory coupling.

## CRediT authorship contribution statement

M.A.L. and D.F.K.: Conceptualization, Methodology. M.A.L., A.S.M., K.V.S., and D.F.K.: Formal analysis, Investigation. M.A.L., N.M.S., A.V.M., and D.F.K.: Writing – Original Draft Preparation. All authors: Writing – Review & Editing.

## Declaration of Generative AI and AI-assisted technologies in the writing process

Artificial intelligence (AI) was used for language editing. No new material was created using AI. The tool employed was Grok (version 3; xAI, Culver City, CA, USA).

## Data availability

The experimental data used for this analysis are available online (https://physionet.org/content/cebsdb/1.0.0/), and software will be made fully available upon the completion of the review process.

